# Molecular phylogenetics and character evolution in *Haplanthodes* (Acanthaceae), an endemic genus from peninsular India

**DOI:** 10.1101/2020.11.08.373605

**Authors:** Siddharthan Surveswaran, Neha Tiwari, Praveen K. Karanth, Pradip V. Deshmukh, Manoj M. Lekhak

## Abstract

*Haplanthodes* (Acanthaceae) is an Indian endemic genus with four species. It is closely related to *Andrographis* which is also mainly distributed in India. *Haplanthodes* differs from *Andrographis* by the presence of cladodes in the inflorescences, sub actinomorphic flowers, stamens included within the corolla tube, pouched stamens and oblate pollen grains. To understand the phylogenetic relationship of *Haplanthodes* with *Andrographis* and *Haplanthus*, another related genus, we used four plastid markers, *matK, rbcL, psbA-trnH* and *trnGR* to construct a molecular phylogeny. Our results established the monophyly of this genus and revealed a sister relationship to *Andrographis* and *Haplanthus*. Further, to understand the historical biogeography of the genus, we inferred the divergence time and performed ancestral area reconstruction. Our analyses suggest that *Haplanthodes* has evolved during Late Miocene 5.85 Ma [95%HPD: 2.18-10.34 Ma] in peninsular India where it might have shared a common ancestor with *Andrographis*. To understand character evolution, the ancestral states of important morphological characters were inferred based on the equal rate model and discussed. The generic status of *Haplanthus* is not resolved due to incomplete sampling.

## Introduction

India is home to about 50 endemic genera of plants, most of which are concentrated in the Western Ghats and the Northeast mountainous region of India (Irwin and Narasimhan 2011; Singh et al 2015). The Western Ghats is one among the 34 global biodiversity hotspots which offer diverse habitats for speciation and contributes to the enormous enrichment of biological diversity in peninsular India (Myers et al 2000; Mittermeier et al 2004). It is a chain of mountains running between the 8°-21°N latitudes running along the West coast of peninsular India. Across taxonomic groups the Western Ghats harbour around 1700 endemic taxa (Mittermeier et al 2004). Endemism and species richness are not uniformly distributed across the Western Ghats. Overall, the southern Western Ghats possess a higher degree of endemism and species richness than the northern and central part of Western Ghats (Pascal 1988; Daniels 1992; Gimaret-Carpentier et al 2003; Davidar et al 2007; Joshi and Karanth 2011; Divya et al 2020).

The Western Ghats has a range of habitats and very diverse forest types such as; deciduous forests, tropical evergreen forests, montane forests and high elevation plateaus. Studies on past climate have shown that the present day diversity can be attributed to recent climatic changes and opening up of new ecological niches. The rise and extension of the great Himalayan range during the middle Miocene (~15 Ma) (Ding et al 2017) might have resulted in monsoon seasonality and increased temperature in peninsular India including the Western Ghats. During the middle Miocene, most of peninsular India was forested (Guleria 1992; Morley 2018) to eventually become more open and arid during late Miocene (11.5 - 5.3 Ma) and Pliocene (1.95 - 5.3 Ma) corresponding to global cooler and drier global climate. For instance, until 8 Ma C3 vegetation dominated in India than during the late Miocene and early Pliocene (8 to 5 Ma) expansion of C4 grasslands initiated and successively dominated the vegetation of India (Cerling et al 1997; Edwards et al 2010). There might have been several floral exchanges from Southeast Asia into peninsular India (Page and Surveswaran 2014; Sen et al 2019).

The study of endemism in the Western Ghats would help us understand the historical biogeography and palaeoclimate underlying the floral assemblages in peninsular India. Most of the endemic genera in India are mostly orchids or grasses (Irwin and Narasimhan 2011). Most of the Indian endemic dicot genera are monotypic whose molecular phylogenetics has not been studied. *Haplanthodes* Kuntze (Acanthaceae Juss.) (Fig. 1) with four species is one such genus in which no phylogenetic evaluation has been done.

**Fig. 1.**
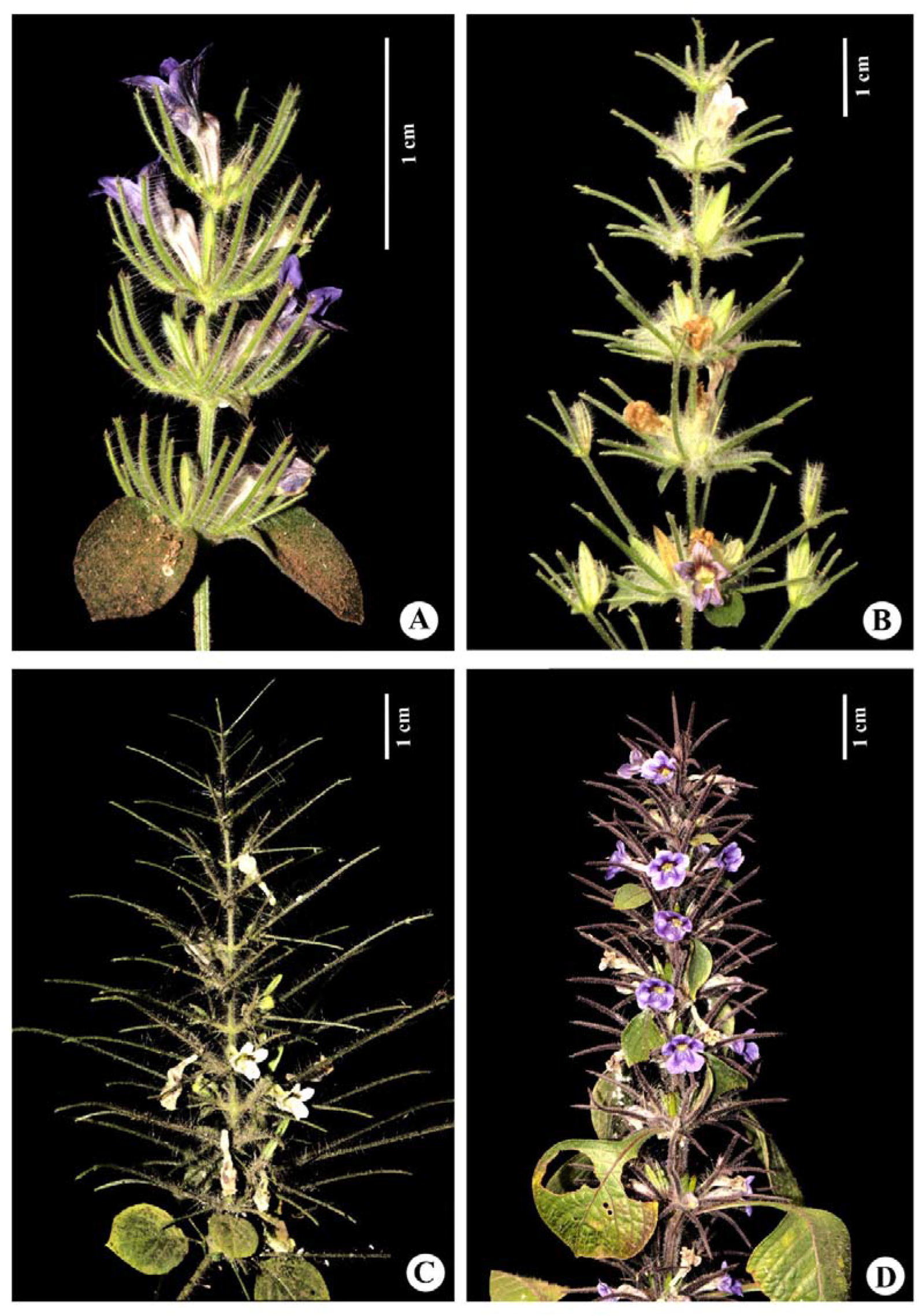
Morphology of the inflorescence of *Haplanthodes* species. A. *H. neilgherryensis*, B. *H. plumosa*, C. *H. tentaculata* and D. *H. verticillata*.

The subfamily Acanthoideae is distributed in the Old World and its tribe Andrographideae is purely Asian. The genera in Andrographideae are *Andrographis* Wall. ex Nees, *Cystacanthus* T. Anderson, *Gymnostachyum* Nees, *Haplanthodes* Kuntze, *Haplanthus* Nees. and *Phlogacanthus* Nees. *Cystacanthus* contains 10 species present in India, China, Thailand, Myanmar and Vietnam. *Gymnostachyum* is present in southeast Asia and India with about 50 species. *Phlogacanthus* contains 43 species abundantly distributed in southeast Asia with a few species in India. *Andrographis* Wall. ex Nees is composed of annual herbaceous plants, native to Asia with 26 species (POWO 2019). Out of the 26 species, 19 species (and 4 varieties) are endemic to India (Singh et al 2015). *A. stellulata* C.B.Clarke, *A. rothii* C.B.Clarke, *A. stenophylla* C.B.Clarke, *A. producta* (C.B.Clarke) Gamble, *A. lobelioides* Wight, *A. lineata* Nees, *A. atropurpurea* (Dennst.) Alston, *A. affinis* Nees, *A. serpyllifolia* (Vahl) Wight, *A. neesiana* Wight and *A. laxiflora* (Blume) Lindau are endemic to the Indian sub-continent. *A. echioides* Nees is present in India, Sri Lanka and Myanmar. *A. elongata* T. Anderson is present in India and Thailand while *A. ceylanica* and *A. alata* Nees are endemic to India and Sri Lanka. *A. paniculata* Nees is the only widely distributed species and occurs in southern and south-eastern Asia (including, India, Sri Lanka, China, Thailand etc.). It is also widely cultivated all over the world due to its high medicinal value. The active component of *A. paniculata* is Andrographolide, a bicyclic diterpenoid lactone. The decoction of *A. paniculata* has proven to be an effective cure for Dengue infection and there are reports which prove its antiviral activity (Panraksa et al 2017). *Indoneesiella* was a genus endemic to India but now its species are subsumed into *Andrographis* (A. echioides (L.) Nees and *A. longipedunculata* (Sreem.) L.H. Cramer) (Cramer 1996; Gnanasekaran et al 2020). *Haplanthus* comprises 4 species including two varieties. Thus, the tribe is Asian with several taxa distributed in the Indian subcontinent.

*Haplanthodes* is endemic to peninsular India, mostly within the Western Ghats. *Haplanthodes* belongs to the family Acanthaceae which comprises 242 genera and 3947 species under 4 subfamilies: Acanthoideae, Avicennioideae, Nelsonioideae and Thunburgioideae (The Plant List 2013). McDade et al. (McDade et al 2008) described an updated backbone phylogeny of Acanthaceae and resolved various subfamilies and tribes. The genus *Haplanthodes* is a member of the subfamily Acanthoideae under the tribe Andrographideae which in turn consists of six genera. It is closely related to *Andrographis* and the recently resurrected genus *Haplanthus* Nees (Gnanasekaran et al 2016) with respect to morphology and biogeography.

The species of *Haplanthodes* are: 1) *H. neilgherryensis* (Wight) R.B.Majumdar 2) *H. plumosa* (T. Anderson) Panigrahi & G.C. Das 3) *H. tentaculata* (L.) R.B.Majumdar and 4) *H. verticillatus* (Roxb.) R.B.Majumdar (Panigrahi and Das, 1981). *H. neilgherryensis* is the most common among the four. The genus is distributed in the states of Goa, Gujarat, Karnataka, Kerala, Madhya Pradesh, Maharashtra, Rajasthan and Tamil Nadu (Fig. 2). All species grow mostly along the edges of the forest, lateritic slopes or bunds, forest paths, plateaus, roadsides and as under shade herbs in deciduous forests. They grow in open spaces, between 204 □ 668 m in elevation.

**Fig. 2.**
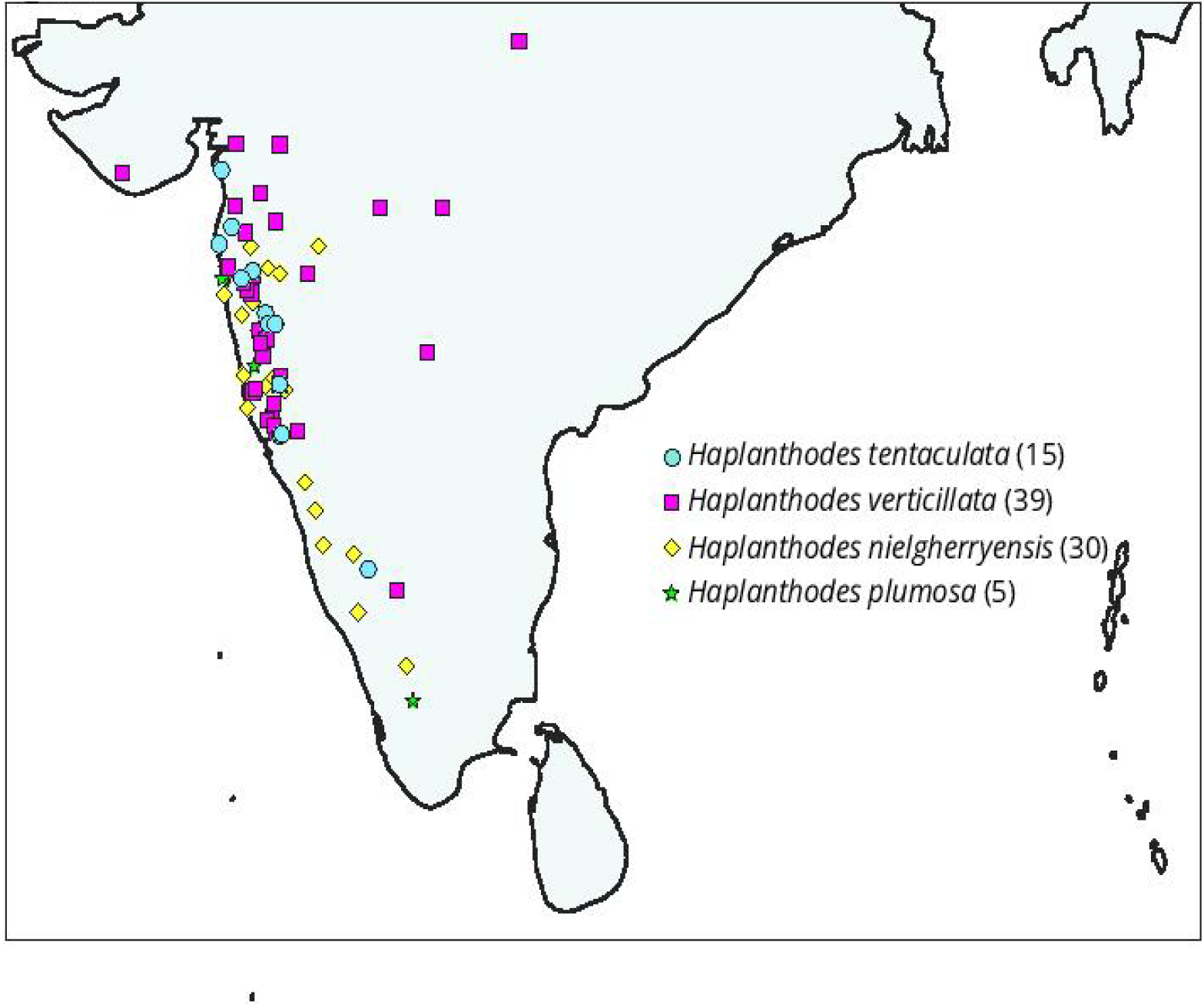
Distribution of *Haplanthodes* species.

*Haplanthodes* has a common biogeographical distribution and some morphological similarities to *Andrographis*. However, it differs from the latter in several morphological characters (Table 1). *Haplanthodes* occupies the arid and semi-arid ecological niches in the Western Ghats. It can be distinguished with the other members of tribe Andrographideae by presence of cladode, densely hairy trichomes and hygroscopic seeds. These characters appear to have independently evolved in this genus and might be an adaptation to its present-day habitat. Other studies also have linked the acquisition of cladode as an adaptation to dry conditions (Fukuda et al 2005; Pimienta□Barrios et al 2005; Nakayama et al 2012). Further, *Haplanthodes* possesses an oblate type of pollen grain shape which has evolved from the prolate type of pollen grain as the ancestral character. Some studies have suggested that the change in shape and size of pollen may be affected by ecological factors such as water availability (Crisp et al 2009)) and a shift in the mode of pollination (Torres 2000).

**Table 1.**
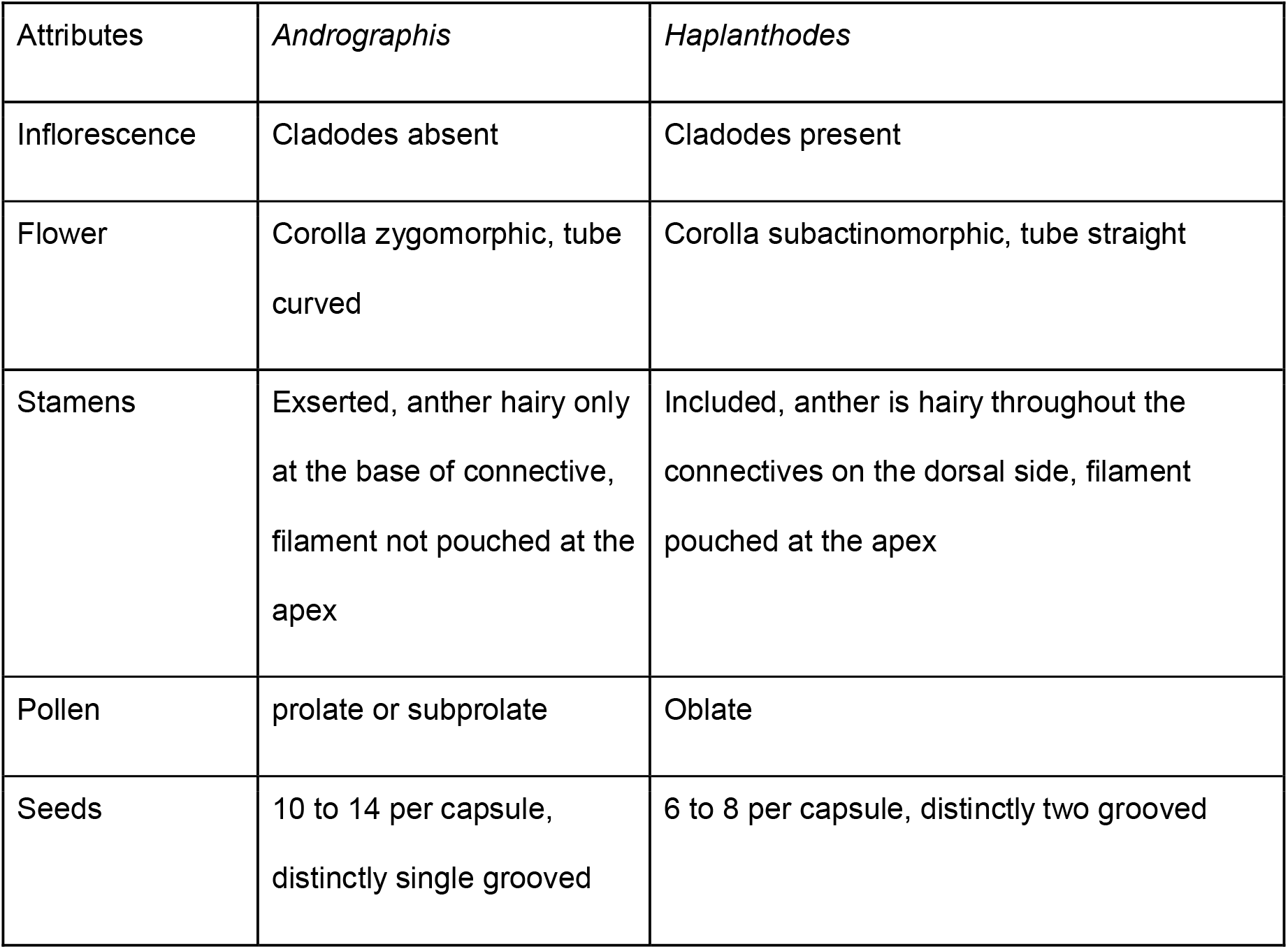
Comparison of morphological characters of *Andrographis* and *Haplanthodes*

| Attributes | *Andrographis* | *Haplanthodes* |
| --- | --- | --- |
| Inflorescence | Cladodes absent | Cladodes present |
| Flower | Corolla zygomorphic, tube curved | Corolla subactinomorphic, tube straight |
| Stamens | Exserted, anther hairy only at the base of connective, filament not pouched at the apex | Included, anther is hairy throughout the connectives on the dorsal side, filament pouched at the apex |
| Pollen | prolate or subprolate | Oblate |
| Seeds | 10 to 14 per capsule, distinctly single grooved | 6 to 8 per capsule, distinctly two grooved |

Given this scenario, we explore the origin of *Haplanthodes* in the Western Ghats. The specific objectives of this project were: 1) to assess the monophyly of *Haplanthodes* and its relationship with putative sister genera *Andrographis* and *Haplanthus*, 2) to elucidate the historical biogeography of *Haplanthodes* along with closely related taxa such as *Andrographis* and *Haplanthus* and 3) to estimate the ancestral character in Andrographidae and evolution of characters in *Haplanthodes* and *Andrographis*. For this, we used four molecular markers from the plastid genome and employed Bayesian phylogenetic reconstruction followed by molecular dating of the clades, biogeography and character evolution analyses.

## Materials and Methods

### Taxon sampling and DNA extraction

Plant tissue samples were collected from various places from peninsular India as listed in Table 2. Our collection includes all described species of genus *Haplanthodes* (including one putative new species), one *Haplanthus* (out of four described) and one *Andrographis* (out of 26 described). The sequences of rest of the tribe Andrographideae, including several other genera of Acanthaceae, were downloaded from the GenBank database (Online Resource 1). A total of 68 new GenBank accessions representing 10 taxa were used in the present study; multiple individuals of each species have been sampled to reduce the chance of sampling biases.

**Table 2.** Taxa included in the study, their locality, voucher information and GenBank accession number of the gene regions used.

| Species | Location | Voucher | *matK* | *rbcL* | *psbA-trnH* | trnGR |
| --- | --- | --- | --- | --- | --- | --- |
| *Haplanthodes neilgherryensis* | Rajapur, Maharashtra | MML-532 | MT445703 | MT445687 | MT445668 | MT445650 |
|  |  | (HN1) |  |  |  |  |
|  | Amboli Ghat,<br>Maharashtra | MML-529<br>(HN2) | MT445<br>704 | MT445<br>688 | MT445<br>669 | MT445<br>651 |
|  | Jamgaon,<br>Karnataka | MML-526<br>(HN3) | MT445<br>705 | - | MT445<br>670 | MT445<br>652 |
|  | Amboli Ghat,<br>Maharashtra | MML-534<br>(HN4) | MT445<br>706 | MT445<br>689 | MT445<br>671 | MT445<br>653 |
|  | B shettigeri<br>(Coorg),<br>Karnataka | HN5 | MT445<br>707 | MT445<br>690 | MT445<br>672 | MT445<br>654 |
| *H. plumosa* | Amboli Ghat,<br>Maharashtra | MML-546<br>(HP1) | MT445<br>708 | MT445<br>691 | MT445<br>673 | MT445<br>655 |
|  | Amboli Ghat,<br>Maharashtra | MML-524<br>(HP2) | MT445<br>709 | MT445<br>692 | MT445<br>674 | MT445<br>656 |
| *H. verticillata* | Panhala,<br>Budhwar<br>peth,<br>Maharashtra | MML-553<br>(HV1) | MT445<br>710 | MT445<br>693 | MT445<br>675 | MT445<br>657 |
|  | Hasane<br>Radhanagari,<br>Maharashtra | MML-554<br>(HV2) | MT445<br>711 | MT445<br>694 | MT445<br>676 | MT445<br>658 |
| *H. tentaculata* | Tillari Ghat,<br>Maharashtra | MML-531<br>(HT1) | MT445<br>712 | MT445<br>695 | MT445<br>677 | MT445<br>659 |
| *Haplanthodes. sp. 1* | Nashik,<br>Maharashtra | MML-544<br>(HS1) | MT445<br>713 | MT445<br>696 | MT445<br>678 | MT445<br>660 |
| *Haplanthus ovatus* | Sita River,<br>Shivamogga<br>district,<br>Karnataka | HJCB-5432 | MT445<br>714 | MT445<br>697 | MT445<br>679 | MT445<br>661 |
|  | Bodamalai<br>Hills, Salem,<br>Tamilnadu | HJCB-1962 | - | MT445<br>698 | MT445<br>680 | MT445<br>662 |
| *Andrographis alata* | Shevaroy<br>Hills, Salem,<br>Tamil Nadu | HJCB-2259 | - | - | MT445<br>681 | MT445<br>663 |
|  | KMTR,<br>Tirunelveli,<br>Tamilnadu | HJCB-2250 | MT445<br>715 | MT445<br>699 | MT445<br>682 | MT445<br>664 |
| *Andrographis* | Talakaveri, | HJCB- | - | - | MT445 | MT445 |
| *megamalayana 1* | Kodagu,<br>Karnataka | 1190 |  |  | 683 | 665 |
| 2 | Talakaveri,<br>Kodagu,<br>Karnataka | HJCB-1454 | MT445<br>716 | MT445<br>700 | MT445<br>684 | - |
| *Andrographis paniculata* | Batrepalli,<br>Anantapur,<br>Andhra<br>Pradesh | HJCB-0572 | MT445<br>717 | MT445<br>701 | MT445<br>685 | MT445<br>666 |
| *Andrographis glandulosa* | Pulivendala<br>ghat, Kadapa<br>district,<br>Andhra<br>Pradesh | HJCB-E-265 | - | MT445<br>702 | MT445<br>686 | MT445<br>667 |

Leaf samples were collected and stored in silica gel before DNA extraction. The total genomic DNA was isolated using the Qiagen DNeasy Plant Mini Kit (Qiagen, Hilden, Germany). Four plastid markers, two coding regions *matK* and *rbcL* and *two* non-coding, *psbA-trnH* and *trn*GR, were used in the study. Forward and reverse primers for *matK, rbcL* and *trn*GR were obtained from (Tripp 2007; Yu et al 2011; Bafeel et al 2011), respectively. Forward and reverse primers for *psb*A-*trn*H were obtained from (Sang et al 1997) and (Tripp 2010), respectively. Primer sequences, product sizes and annealing temperature information are listed in Online Resource 2.

Each 25 μL polymerase chain reaction (PCR) mixture consisted of 2.5 μL of 10x NEB Taq Buffer, 1.5 μL of 2.5 mM MgCl_2_, 1.5 μL of 10 mM dNTPs mix,1.5 U of NEB Taq DNA polymerase (New England Biolabs, USA) and 10 pmol of each primer and 10-200 ng of DNA template. The PCR conditions were as follows: initial denaturation at 94°C for 2 min, followed by 40 cycles each of denaturation at 94° C for 30 sec, annealing at 48° C for 30 sec, extension at 68° C for 1 min and a final extension at 68 °C for 10 min. The annealing temperature was 48 °C for *matK* while it 52 °C for *rbcL* and *psbA-trnH* and 54 °C for *trn*GR (Online Resource 1). PCR products were visualized on 1% agarose gels stained with ethidium bromide. PCR product purification and Sanger sequencing were done at Barcode Biosciences, Bangalore, India.

### Sequence analysis and alignment

Sequence quality was checked using Chromas v 2.6.6 (Technelysium, www.technelysium.com.au). Sequences were aligned using the MUSCLE algorithm as implemented in MEGA v 7.0 (Kumar et al 2016) and manually checked to avoid any artefacts using AliView ver. 1.17 (Larsson 2014). The length of the individual genes were 729, 498, 760 and 812 base pairs (bp) for *matK, rbcL, psbA-trnH* and *trnGR*, respectively. The number of sequences, parsimoniously informative characters (PIC), etc. are presented in Online Resource 3.

### Phylogenetic Analysis

Since all the markers were from the plastid genome, combined analysis of the four markers was performed. The concatenated supermatrix of the four markers consisted of 2799 aligned characters. Partitionfinder v 2.1.1 (Lanfear et al 2017) was used to find the best model of sequence evolution and the partitioning scheme using the Bayesian information criterion (BIC). Partitions of data and best-fitting models are provided in Online Resource 4.

Maximum likelihood (ML) analysis was performed using RAxML HPC v. 8.2.12 (Stamatakis 2014) as implemented in CIPRES science gateway (Miller et al 2010). Ten independent runs were initiated, along with 1000 rapid bootstrap replicates to estimate the node support. For Bayesian analysis, a nexus file was prepared to contain the alignment data set and MrBayes block, and the analysis was carried out using MrBayes v 3.2.7a (Ronquist et al 2012) on the CIPRES portal (http://www.phylo.org/). Two independent runs each consisting of two of Markov Chain Monte Carlo (MCMC) chains were run for 20 million generations with sampling every 2000 generations. The chain convergence across runs was determined using the average standard deviation of split frequency < 0.01. The software Tracer v1.7.1 (Rambaut et al 2018) was used to plot the log-likelihood values of two runs and to check the stationarity phase using the expected sample size (ESS) > 200. The consensus tree was summarized post burn-in of initial 25% sampled trees. Trees were visualized in FigTree v1.4.4 (Rambaut 2018).

### Divergence time estimation

A dataset containing 71 taxa encompassing tribes, Andrographidae, Barlerieae, Whilfieldieae, Ruelliae and Justicieae were compiled from newly sequenced data and GenBank data. *Martynia* and *Sesamum* were used as outgroups in the Bayesian analysis (Tripp and McDade 2014). The dataset consisted of 2784 of which 1129 were variable and among those 716 were parsimoniously informative. The best model of sequence evolution as selected by the Akaike Information Criterion (AIC) was GTR + I + Γ as determined using Partitionfinder v2.1.1 (Lanfear et al 2017). Dating of the nodes was inferred using an uncorrelated relaxed clock model as implemented in BEAST v1.8.4 (Drummond and Rambaut 2007; Suchard and Rambaut 2009). For the dating of the nodes using fossil information, two calibrations were picked and implemented as described in (Tripp and McDade 2014). One of the fossils was the Miocene age of the crown of Barlerieae from (Kuyl et al 1955). The second calibration point was 10-12 Ma as characterized by fossil pollen similar to modern *Bravaisia (Graham 1976)*. The fossil ages were modelled with the lognormal distribution as described in (Tripp and McDade 2014). The crown age of Barlerieae was assigned a log mean of 5.5, log standard deviation (stdev) of 1.1 and an offset of 5.3 Ma under a lognormal distribution with the option of sampling the mean from real space in the distribution. The interval between 95% and 5% intervals were 23.64 and 5.792 Ma, respectively which roughly encompasses the Miocene epoch. The age of the clade consisting of *Sanchezia* Ruiz & Pav. and *Strobilanthes* Blume (Justiceae) was assigned a log mean of 1, stdev of 0.5 and an offset of 10 Ma. The sampling was set to be under the real space and the 95% to 5 % intervals of the lognormal distribution were 12.01 and 10.39 Ma. Hundred million runs of the MCMC were performed with sampling every 10000 runs. The BEAST analysis was also carried out in the CIPRES portal (www.phylo.org). The maximum clade credibility tree was recovered in Treeannotator v1.8.4 (Drummond et al 2012) and visualized in FigTree v1.4.4.

### Morphological Character Evolution

To understand the evolution of diagnostic morphological characters in *Haplanthodes* we assessed four binary and four multistate traits which were cladode (presence or absence), the shape of corolla: (actinomorphic or subactinomorphic), corolla tube (straight or curved), stamens (exserted or included), anthers (bearded or glabrous or hairy only on the dorsal side), filaments (pouched or not), pollen shape (prolate or oblate), seeds shape (distinctly single grooved, two grooved or not grooved). The character coded data matrix is provided (Online Resource 5).

The time tree from above divergence time estimation analysis was used for ancestral state reconstruction and biogeographical analysis. The dataset used was pruned to 62 taxa where multiple individuals of the same species were removed. The BEAST maximum clade credibility tree obtained was pruned to represent only the tribe Andrographidae. We used ancestral state reconstruction as implemented in “rayDISC” function for discrete data in the corHMM package (Beaulieu et al 2013; R Core Team 2019) in R version 3.6.0 (R Core Team 2019). rayDISC assumes a constant rate of evolution across all branches and calculates the probability of the ancestral character at each node. For each binary trait, we used the equal rate (ER) model (Mk model) which assumes a single rate of transition among all possible states. Additionally, we used the all rates different (ARD) (AssymMK model, a rate heterogeneity model, (Pagel 1999)) which allows different rates for each transition between two states. The best fit of the two models was selected using the Akaike Information Criterion (AIC) and the log-likelihood values were computed from the corHMM package.

### Ancestral area reconstruction

Ancestral area reconstruction was performed based on the dispersal, extinction and cladogenesis (DEC) model using the package LAGRANGE v 20130526 (Ree and Smith 2008). Andrographideae is an Old World tribe and only two major distributional areas namely, tropical South East Asia and tropical South Asia (peninsular India). These two regions were coded as A (South East Asia) and B (peninsular India). The pruned ultrametric tree for the character mapping comprising genera of Andrographideae was used for ancestral area reconstruction. The maximum ranges allowed were two (A, B) and both areas were allowed to be combined. The rates of dispersal were allowed to be equal (symmetric) between the two areas across the entire time frame because these are landmasses adjacent to each other.

## Results

The dataset for the combined analysis of the four plastid loci consisted of 70 taxa and 2799 aligned characters. The Bayesian phylogenetic inference tree (Fig. 3) was similar in topology with the ML tree topology (data not shown). The genus *Haplanthodes* along with other closely related genera, *Andrographis* and *Haplanthus* formed a well-supported clade. *Haplanthus* was found nested within *Andrographis*, with *A. alata* as a sister. However, this relationship was not statistically well supported. *Andrographis echioides* (syn. *Indoneesiella echioides*) was nested in the genus *Andrographis* indicating that the genus name *Indoneesiella* can be subsumed into *Andrographis*. Tribes Andrographideae, Barlerieae, Whitfieldieae, Justicieae and Ruellieae of the subfamily Acanthoideae were monophyletic with high statistical support (except for Whitfieldieae).

**Fig. 3.**
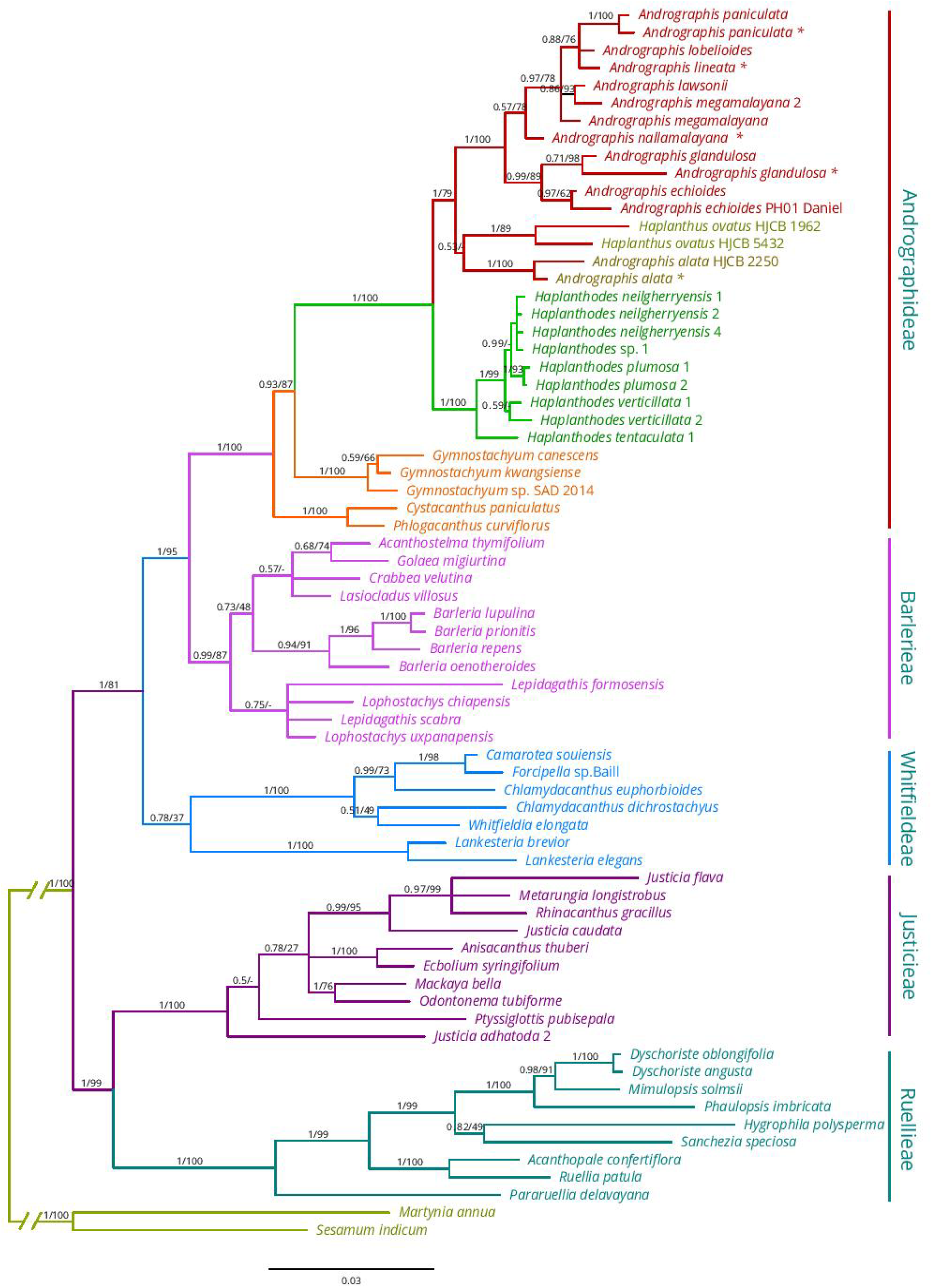
Bayesian phylogenetic tree for *Haplanthodes* including several genera of Acanthaceae. Numbers on branches indicate Bayesian posterior probability (BPP) 0.0 to 1.0 and maximum-likelihood bootstrap support percentages. Scale bar indicates the estimated number of nucleotide substitutions per site.

The molecular dating dataset consisted of 71 taxa and 2784 aligned characters after removal of ambiguously aligned regions. The BEAST dated phylogeny inferred the median age of diversification of *Haplanthodes* as 5.85 Ma, varying between 2.18 to 10.34 Ma (95% highest posterior density (HPD) (Fig. 4, Online Resource 6). The age of the tribe Andrographidae was inferred to be 18.53 Ma (9.6 to 28.02 Ma 95% HPD). The age of family Acanthaceae inferred from our analysis was 36.67 Ma (21.7 to 53.61 Ma 95% HPD). A total of five accessions of *H. neilgherryensis* was used in our analysis and two accessions HN3 and HN5 were observed as a separate clade showing a closer relationship with *H. verticillata*. This was observed in the Bayesian analysis (Fig 3) also (not shown) and these taxa were removed from the analysis. A putative new species (unpublished, voucher MML-54), was found to be nested within the clade of *Haplanthodes* sister to *H. neilgherryensis*.

**Fig. 4.**
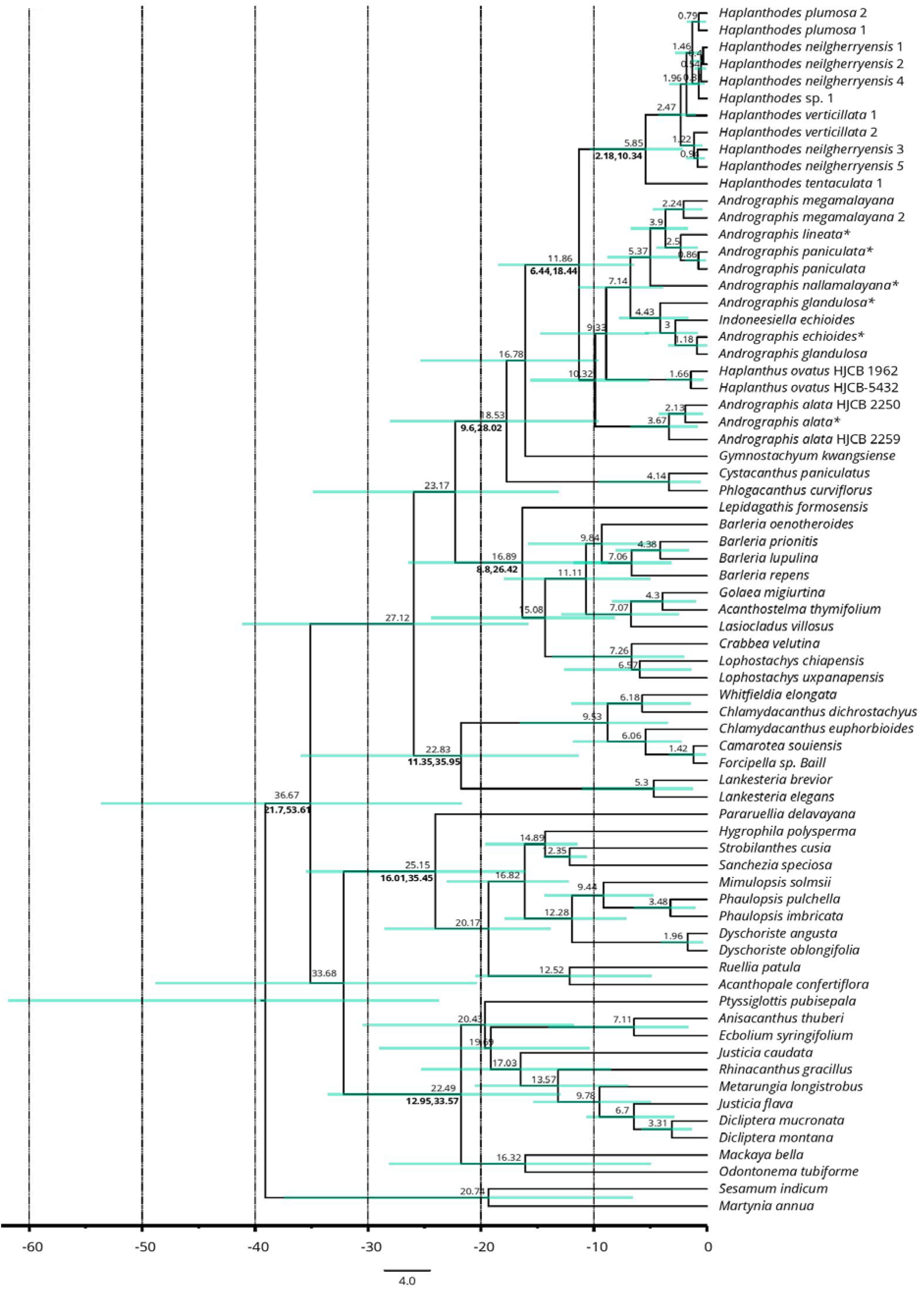
BEAST phylogeny of the subfamily Acanthoideae. The scale beneath the tree indicates the average number of substitutions per unit time.

In the biogeographic analysis, the ancestral area for *Haplanthodes* and *Andrographis* was found to be peninsular India (region B) (Fig. 5). However, the ancestral area of the lineage containing *Haplanthodes, Andrographis* and *Gymnostachyum* was found to be both South East Asia and peninsular India (region AB).

**Fig. 5.**
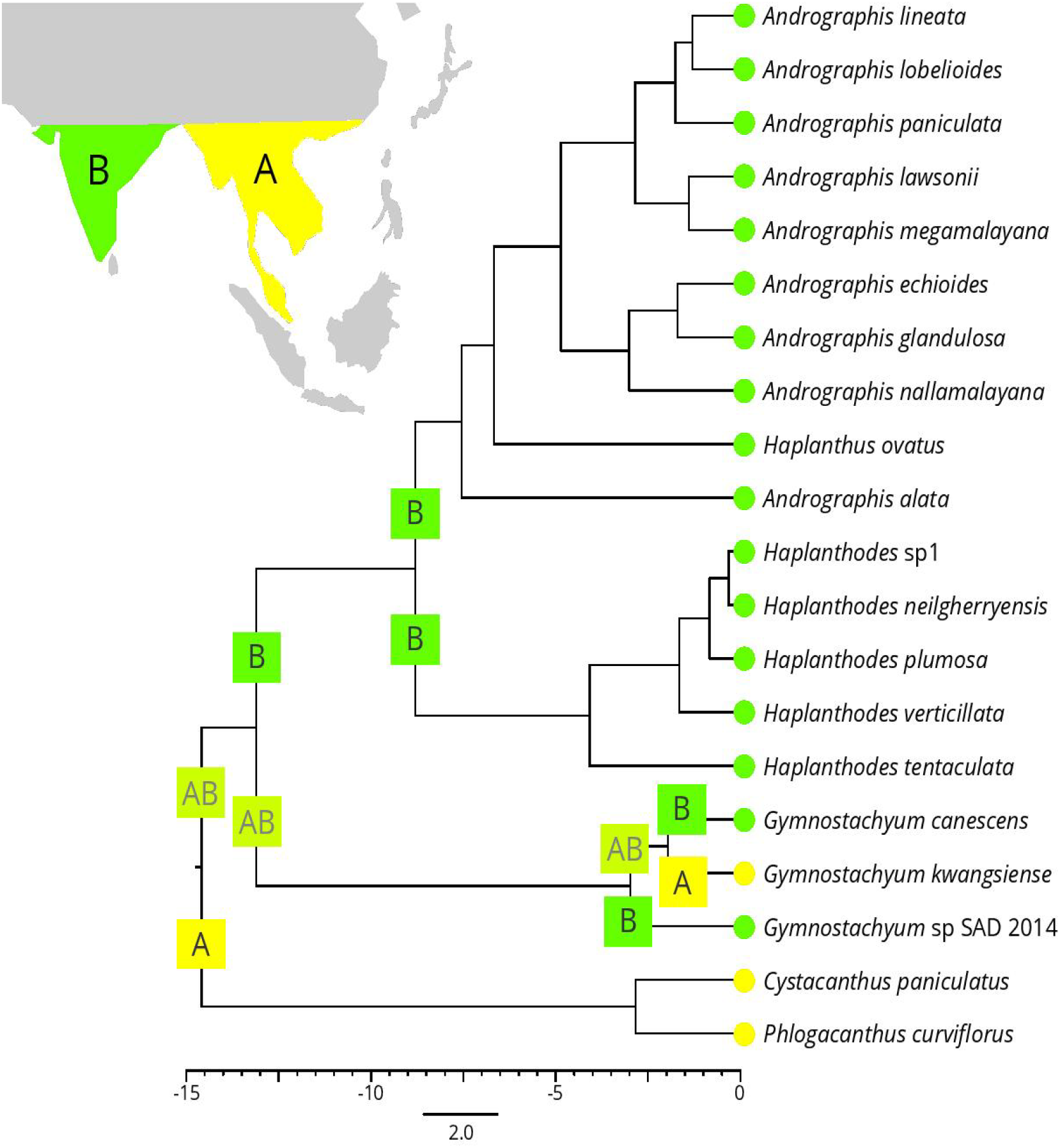
Biogeographical analysis of the genus *Haplanthodes* with respect to the other genera in the tribe Andrographideae using the DEC model as implemented in LAGRANGE based on BEAST derived chronogram. The inferred ancestral regions are indicated on the upper and lower portions of the split on each node. Only splits within 2 log-likelihood units of the maximum for each node are shown.

A total of ten characters were taken for ancestral state reconstruction analyses. Character evolution was performed using ER and ARD models and the best-fit model was chosen according to the likelihood scores and the AIC (Online Resource 7). The ER model was the best-fit model for character evolution and is indicated in Figs. 5, 6, and 7. The ARD model analysis for the study of ancestral character evolution is available in Online Resource 8. The presence of the cladode is an important apomorphy for *Haplanthodes* and the ancestral character was inferred to be the absence of cladode (Fig. 6A). Similarly, the flower shape namely subactinomorphic is homoplasious (convergent) for *Haplanthodes* and *Haplanthus* and it seems to have evolved from genera having zygomorphy as ancestral (Fig. 6B). The corolla tube being curved is the ancestral character for the tribe Andrographideae including *Haplanthodes* and the straight corolla tube was found to have evolved from curved corolla tube as seen in *Andrographis* (Fig. 6C). Stamen included inside the corolla tube is a homoplasious character in *Haplanthodes* and *Haplanthus* which might have arisen from exserted stamens (Fig. 6D). The presence of hair on the dorsal side of the anthers is a homoplasy for *Haplanthodes* and *Haplanthus* whereas the ancestral character was inferred to be the bearded type found in *Andrographis* (Fig. 7E). The bearded type where there is a tuft of hairs at the base of the anthers at the point of attachment of filament is symplesiomorphy for *Andrographis* which has been lost in *A. lawsonii*. Similarly, the presence of a pouch at the filament is a homoplasy for *Haplanthodes* and *Haplanthus* both derived from non-pouched filaments (Fig. 7F). Oblate (flattened at the poles) pollen is homoplasy for *Haplanthodes* and *Haplanthus* which have been derived from prolate pollen (Fig. 7G). A seed with two grooves along the sides is an apomorphy in *Haplanthodes* derived from single grooved seeds which are seen in *Andrographis* (Fig. 7H). Presence of hairs in seeds is the ancestral character (plesiomorphy) as seen in *Haplanthodes* whereas glabrous seeds are synapomorphic in *Andrographis* and *Haplanthus* (Fig. 8I). Compressed seeds are apomorphic for *Haplanthus* whereas the non-compressed inflated seeds are the ancestral character as seen in *Haplanthodes* and *Andrographis* (Fig. 8J).

**Fig. 6.**
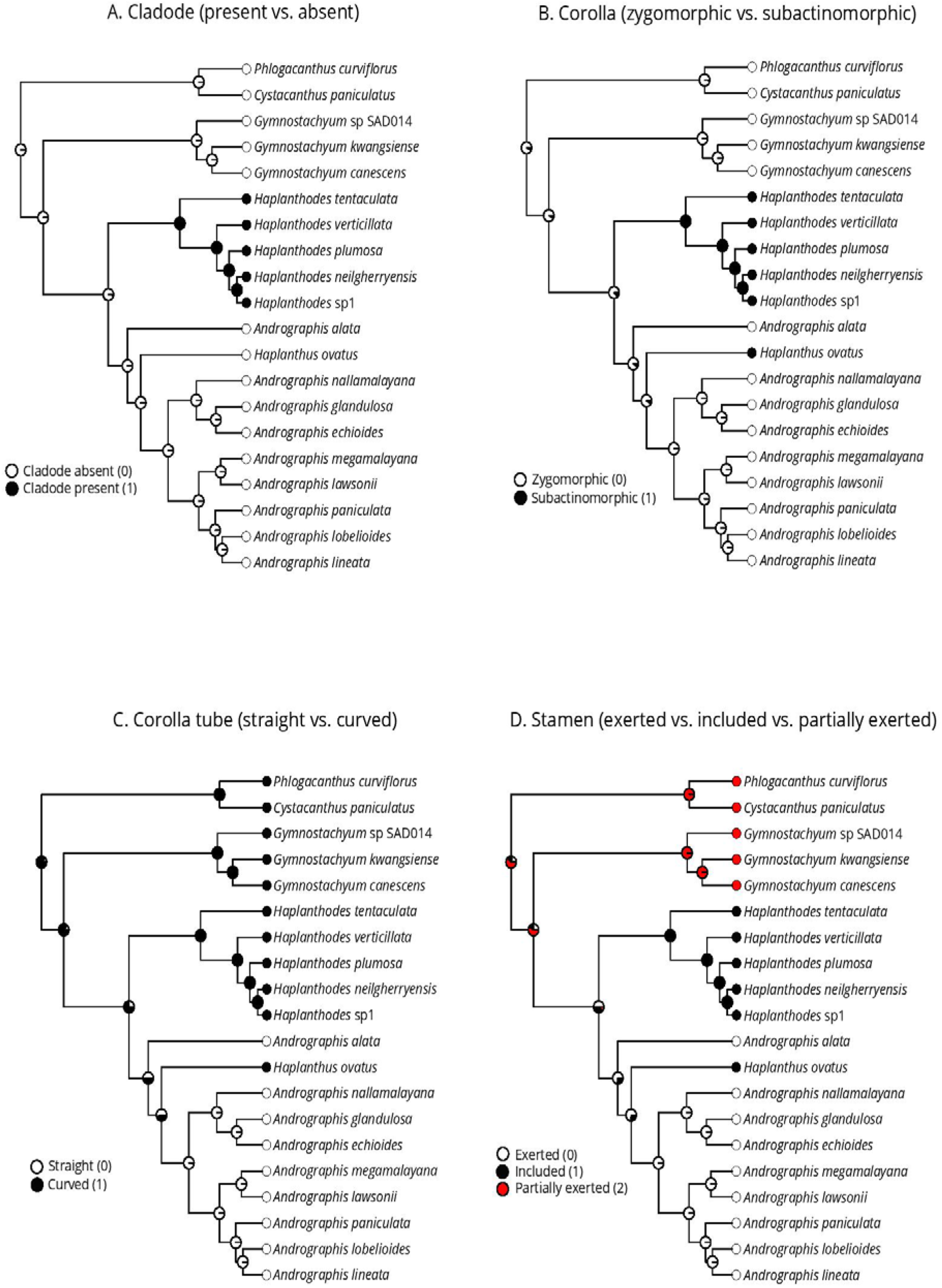
Character evolution in *Haplanthodes*, Part I. Circles at nodes reflect ancestral character states (see the key to codes on each figure) with sizes of differently coloured wedges indicating the likelihood of the presence of each state at that node. A, Cladodes: absent vs. present; B, Corolla shape: zygomorphic vs. subactinomorphic; C, Corolla tube: straight vs. curved; D, Stamen: exserted vs. inserted.

**Fig. 7.**
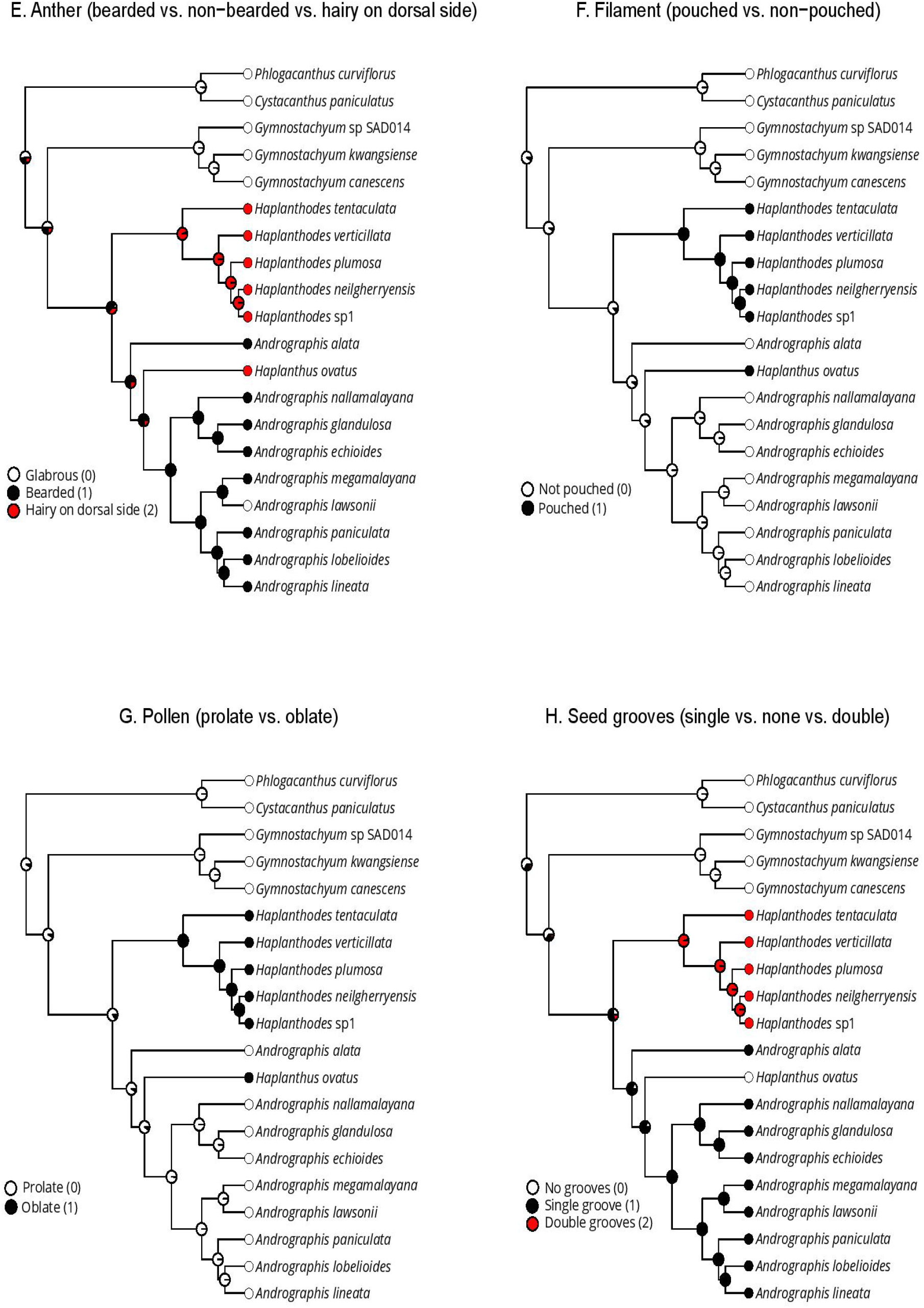
Character evolution in *Haplanthodes*, Part II. Circles at nodes reflect ancestral character states (see the key to codes on each figure) with sizes of differently coloured wedges indicating the likelihood of the presence of each state at that node. E, Anther: bearded vs. non-bearded; F, Filament: pouched vs. not pouched; G, Pollen shape: prolate vs. oblate; H, Seed grooves: none vs. single vs. double.

**Fig. 8.**
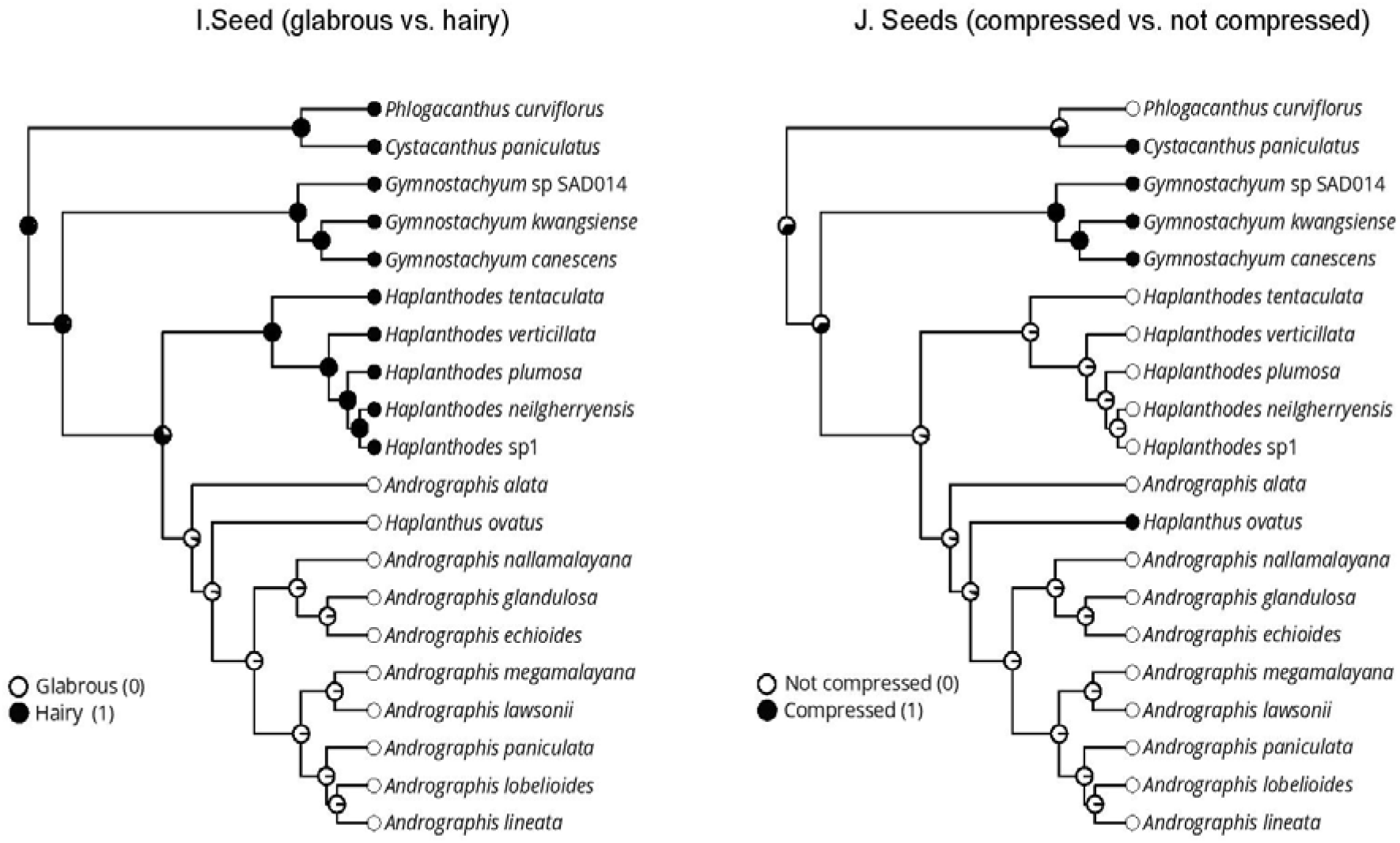
Character evolution in *Haplanthodes*, Part III. Circles at nodes reflect ancestral character states (see the key to codes on each figure) with sizes of differently coloured wedges indicating the likelihood of the presence of each state at that node. I, Seed: glabrous vs. hairy; J, Seed shape: compressed vs. not compressed.

## Discussion

### Taxonomic conflicts between the sister genera of *Haplanthodes*

In the current study, four species of the genus *Haplanthodes (H. neilgherryensis, H. plumosa, H. tentaculata* and *H. verticillata)*, and one species of the genus *Haplanthus (H. ovatus)* were analyzed in conjunction with other genera of the tribe Andrographideae. Our results confirm that the genus *Haplanthodes* is reciprocally monophyletic with respect to *Andrographis* warranting a separate generic status. In the study by Gnanasekaran et al (Gnanasekaran et al 2016), *Haplanthus* was resurrected as a genus from *Andrographis*, based on several morphological characters. While *Haplanthodes* is endemic to peninsular India, the four species of *Haplanthus* are distributed both in peninsular India as well as southeast Asia. One species of *Haplanthus, H. ovata*, was used in our study and it was found nested within genus *Andrographis* (Fig 3) without any scope for taxonomic resolution. However, we are confident that better sampling in future studies will elucidate the clear taxonomic status of *Haplanthus*. The absence of cladode and glabrous seeds are synapomorphies for *Andrographis* and *Haplanthus. Andrographis*, a predominantly Indian genus is paraphyletic with *Haplanthus* but this needs to be confirmed with a complete sampling of *Haplanthus*. The generic status of *Indoneesiella* is not valid as it is nested within the clade of *Andrographis* with closest molecular phylogenetic relationship to *Andrographis glandulosa* supporting morphological data (Cramer 1996).

### Putative species complexes

We observed that two accessions of *H. nielgherryensis*, HN3 and HN5 were observed as a separate clade showing a closer relationship with *H. verticillata* (Fig 4). A putative new species which is morphologically distinct characters (unpublished, voucher MML-54) was found closely nested with *H. nielgherryensis* (Fig 3, 4). These observations suggest that there might be cryptic species within this taxon and more markers and dense population-level markers should be used to resolve this issue.

### Origin and diversification of *Haplanthodes*

(Tripp and McDade 2014) published a dated phylogeny of Acanthaceae *s.l*. using different fossil calibrations and maximum age constraints. The two fossil calibrations and the modelling of the age distribution in our study were from the above paper without maximum age constraint. The age of the Acanthaceae s.s. inferred from our study was 36.67 Ma (53.61 - 21.7 Ma 95% HPD) which is slightly younger than their inference with 4 fossil constraints and without maximum age enforced (Table 4, analysis 2a), 50.7 Ma (43.2-60.3 95% HPD). However, the age of the tribe Andrographideae was inferred to be 18.53 Ma (28.02 - 9.6 95% HPD) which is much closer to their inference, 19.5 Ma (28.9–17.3) (Analysis 2a).

The result from our time-calibrated phylogeny suggests the age of *Andrographis* clade as 10.32 Ma, and the age of *Haplanthodes* clade as 5.85 Ma (2.18-10.34 Ma 95% HPD), However, the split between the *Haplanthodes* and *Andrographis* clades has happened at 11.96 Ma (6.44-18.44 Ma 95% HPD). According to the inferred dates, both these genera have diversified during the middle to late Miocene and early Pliocene. This time period corresponds well with the past climatic conditions of aridification and seasonality in peninsular India (Zhisheng et al 2001). During the mid-Miocene, the uplift of the Himalayan mountain range accorded with Miocene thermal maxima might have facilitated the dispersal route for the seasonal evergreen forest elements into India (Morley 2018). So far this hypothesis has been studied and in other groups such as reptiles, freshwater fishes, mammals (Morley 2000; Jacques et al 2015; Klaus et al 2016).

During the mid-Miocene, the Indian monsoon was gaining its seasonality and climatic condition was becoming drier than ever (Ding et al 2017). This might have facilitated the origin, evolution and dispersal of new peninsular Indian endemic taxa including *Haplanthodes. Andrographis*, and *Haplanthodes* grows on open lands, plateaus, and semi-deciduous forest and survives in semi-arid habitat and drier climatic conditions. The result of our divergence estimates the age of genus *Haplanthodes and Andrographis* between late Miocene to early Pliocene might be a result of this aridification and opening up of new habitats. It can be argued that the presence of cladodes, presence of dense trichomes in the plant body, densely hairy hygroscopic seeds and prolate pollen are an indication of adaptation and subsequent diversification to xeric conditions by *Haplanthodes (Crisp et al 2009)*.

The speciation in *Andrographis* and *Haplanthodes* might have taken place in peninsular India as indicated by our ancestral area reconstruction. However, the distribution of *Haplanthus* in south-east Asia suggests that might be dispersal into south-east from peninsular India. However, this hypothesis needs to be tested once all the species of *Haplanthus* are sampled for phylogeny reconstruction.

### Character evolution

From the analysis of character evolution across *Haplanthodes*, we have observed various patterns. First, our study suggests that the cladode is the character which is only present in genus *Haplanthodes*, and evolved independently. A similar hypothesis emerges from our analysis of corolla, pollen shape, filament, anther and seed groves: sub-actinomorphic flower, oblate pollen shape, pouched stamen, hairy dorsal side anther and double grooved seeds evolved independently in *Haplanthodes* and *Haplanthus* from actinomorphic flowers, prolate pollen, non-pouched filament and bearded anther, and single grooved seeds. Regarding seed hair and corolla tube, our results infer that curved corolla tube and hairy seeds were the ancestral characters of *Haplanthodes* and continue further. One other interesting pattern that has emerged from character evolution analysis is that *Haplanthus* shares six common characters with *Haplantodes* which might be due to convergence. Furthermore, *Haplanthus* and *Andrographis* only shared two synapomorphic characters which are the absence of cladode and glabrous seed. However, in phylogenetic analysis, *Haplathus* is nested within the clade of *Andrographis* and grouped together with *Andrographis alata*.

## Conclusion

In conclusion, based on molecular and morphological analysis, our study confirms: i) monophyly of the genus *Haplanthodes*, ii) character evolution analysis unravels that few characters have been evolved independently across the genus *Haplanthodes* which make it suitable to survive in arid conditions iii) due to paucity of molecular, and morphological data, we are uncertain about the separate generic status of *Haplanthus*, although it could be species complex iv) the generic position *Indoneesiella* is not supported by molecular evidence as with previous morphological studies.

## Acknowledgements

SS thanks CHRIST (Deemed to be University) minor research project grant number MIRPDSC-1903 for funding this project. We thank K. Raja Kullayiswamy for providing some specimens for this work and Sandeep Sen for providing his valuable comments on the Manuscript.

## References

Bafeel SO, Arif IA, Bakir MA, et al (2011) Comparative evaluation of PCR success with universal primers of maturase K *(matK)* and ribulose-1,5-bisphosphate carboxylase oxygenase large subunit (rbcL) for barcoding of some arid plants. Plant Omics 4:195–198.

Beaulieu JM, O’Meara BC, Donoghue MJ (2013) Identifying hidden rate changes in the evolution of a binary morphological character: the evolution of plant habit in campanulid angiosperms. Syst Biol 62:725–737. doi: 10.1093/sysbio/syt034

Cerling TE, Harris JM, MacFadden BJ, et al (1997) Global vegetation change through the Miocene/Pliocene boundary. Nature 389:153–158. doi: 10.1038/38229

Cramer LH (1996) Notes on Sri Lankan Acanthaceae. Kew Bulletin 51:553. doi: 10.2307/4117032

Crisp MD, Arroyo MTK, Cook LG, et al (2009) Phylogenetic biome conservatism on a global scale. Nature 458:754–756. doi: 10.1038/nature07764

Daniels RR (1992) Geographical distribution patterns of amphibians in the Western Ghats, India. Journal of Biogeography 521–529.

Davidar P, Rajagopal B, Mohandass D, et al (2007) The effect of climatic gradients, topographic variation and species traits on the beta diversity of rain forest trees. Global Ecol Biogeography 16:510–518. doi: 10.1111/j.1466-8238.2007.00307.x

Ding L, Spicer RA, Yang J, et al (2017) Quantifying the rise of the Himalaya orogen and implications for the South Asian monsoon. Geology 45:215–218. doi: 10.1130/G38583.1

Divya R, Ramesh BR, Karanth PK (2020) Contrasting patterns of phylogenetic diversity across climatic zones of Western Ghats: A biodiversity hotspot in peninsular India. J Syst Evol

Drummond AJ, Rambaut A (2007) BEAST: Bayesian evolutionary analysis by sampling trees. BMC Evol Biol 7:214. doi: 10.1186/1471-2148-7-214

Drummond AJ, Suchard MA, Xie D, Rambaut A (2012) Bayesian phylogenetics with BEAUti and the BEAST 1.7. Mol Biol Evol 29:1969–1973. doi: 10.1093/molbev/mss075

Edwards EJ, Osborne CP, Strömberg CAE, et al (2010) The origins of C4 grasslands: integrating evolutionary and ecosystem science. Science 328:587–591. doi: 10.1126/science.1177216

Fukuda T, Ashizawa H, Suzuki R, et al (2005) Molecular phylogeny of the genus Asparagus (Asparagaceae) inferred from plastid petB intron and petD-rpoA intergenic spacer sequences. Plant Species Biol 20:121–132. doi: 10.1111/j.1442-1984.2005.00131.x

Gimaret-Carpentier C, Dray S, Pascal JP (2003) Broad-scale biodiversity pattern of the endemic tree flora of the Western Ghats (India) using canonical correlation analysis of herbarium records. Ecography 26:429–444.

Gnanasekaran G, Arisdason W, Deng Y (2020) An overview on the validation of the name *Andrographis longipedunculata* (Acanthaceae). Phytotaxa 433:75–76. doi: 10.11646/phytotaxa.433.1.8

Gnanasekaran G, Murthy GVS, Deng YF (2016) Resurrection of the genus *Haplanthus* (Acanthaceae: Andrographinae). Blum - J Plant Tax and Plant Geog 61:165–169. doi: 10.3767/000651916X693185

Graham A (1976) Studies in neotropical paleobotany. II. the Miocene communities of Veracruz, Mexico. Ann Mo Bot Gard 63:787. doi: 10.2307/2395250

Guleria JS (1992) Neogene vegetation of peninsular India. Palaeobotanist 40:285–311.

Irwin SJ, Narasimhan D (2011) Endemic genera of Angiosperms in India: A Review. Rheedea 21:87–105.

Jacques FMB, Shi G, Su T, Zhou Z (2015) A tropical forest of the middle Miocene of Fujian (SE China) reveals Sino-Indian biogeographic affinities. Rev Palaeobot Palynol 216:76–91. doi: 10.1016/j.revpalbo.2015.02.001

Joshi J, Karanth KP (2011) Cretaceous-Tertiary diversification among select Scolopendrid centipedes of South India. Mol Phylogenet Evol 60:287–294. doi: 10.1016/j.ympev.2011.04.024

Klaus S, Morley RJ, Plath M, et al (2016) Biotic interchange between the Indian subcontinent and mainland Asia through time. Nat Commun 7:12132. doi: 10.1038/ncomms12132

Kumar S, Stecher G, Tamura K (2016) MEGA7: molecular evolutionary genetics analysis version 7.0 for bigger datasets. Mol Biol Evol 33:1870–1874. doi: 10.1093/molbev/msw054

Kuyl OS, Muller J, Waterbolk HT (1955) The application of palynology to oil geology, with special reference to western Venezuela. Geol Mijnbouw 17:49–76.

Lanfear R, Frandsen PB, Wright AM, et al (2017) Partitionfinder 2: new methods for selecting partitioned models of evolution for molecular and morphological phylogenetic analyses. Mol Biol Evol 34:772–773. doi: 10.1093/molbev/msw260

Larsson A (2014) AliView: a fast and lightweight alignment viewer and editor for large datasets. Bioinformatics 30:3276–3278. doi: 10.1093/bioinformatics/btu531

McDade LA, Daniel TF, Kiel CA (2008) Toward a comprehensive understanding of phylogenetic relationships among lineages of Acanthaceae s.l. (Lamiales). Am J Bot 95:1136–1152. doi: 10.3732/ajb.0800096

Miller MA, Pfeiffer W, Schwartz T (2010) Creating the CIPRES Science Gateway for inference of large phylogenetic trees. 2010 Gateway Computing Environments Workshop (GCE). IEEE, pp 1–8

Mittermeier RA, Gil PR, Hoffmann M, et al (2004) Hotspots revisited: Earth’s biologically richest and most endangered terrestrial ecoregions Mexico City. Mexico: Conservation International in association with CEMEX.[Google Scholar]

Morley RJ (2018) Assembly and division of the South and South-East Asian flora in relation to tectonics and climate change. Journal of Tropical Ecology 34:209–234.

Morley RJ (2000) Origin and evolution of tropical rain forests. Wiley

Myers N, Mittermeier RA, Mittermeier CG, et al (2000) Biodiversity hotspots for conservation priorities. Nature 403:853–858. doi: 10.1038/35002501

Nakayama H, Yamaguchi T, Tsukaya H (2012) Acquisition and diversification of cladodes: leaf-like organs in the genus Asparagus. Plant Cell 24:929–940. doi: 10.1105/tpc.111.092924

Pagel M (1999) Inferring the historical patterns of biological evolution. Nature 401:877–884. doi: 10.1038/44766

Page NV, Surveswaran S (2014) *Friesodielsia sahyadrica* (Annonaceae), a peculiar new species from the Western Ghats, India. Phytotaxa 158:275. doi: 10.11646/phytotaxa.158.3.7

Panraksa P, Ramphan S, Khongwichit S, Smith DR (2017) Activity of andrographolide against dengue virus. Antiviral Res 139:69–78. doi: 10.1016/j.antiviral.2016.12.014

Pascal J-P (1988) Wet evergreen forests of the Western Ghats of India. Institut francais de Pondichery, Pondicherry, India

Pimienta□Barrios E, Zañudo□Hernandez J, Nobel PS (2005) Effects of Young Cladodes on the Gas Exchange of Basal Cladodes of *Opuntia ficus□indica* (Cactaceae) under Wet and Dry Conditions. Int J Plant Sci 166:961–968. doi: 10.1086/449317

POWO (2019) Plants of the World Online. Facilitated by the Royal Botanic Gardens, Kew. http://www.plantsoftheworldonline.org/. Accessed 22 Jul 2020

Rambaut A (2018) FigTree v1.4. https://github.com/rambaut/figtree. Accessed 12 May 2020

Rambaut A, Drummond AJ, Xie D, et al (2018) Posterior summarization in bayesian phylogenetics using tracer 1.7. Syst Biol 67:901–904. doi: 10.1093/sysbio/syy032

Ree RH, Smith SA (2008) Maximum likelihood inference of geographic range evolution by dispersal, local extinction, and cladogenesis. Syst Biol 57:4–14. doi: 10.1080/10635150701883881

Ronquist F, Teslenko M, van der Mark P, et al (2012) MrBayes 3.2: efficient Bayesian phylogenetic inference and model choice across a large model space. Syst Biol 61:539–542. doi: 10.1093/sysbio/sys029

R Core Team (2019) R: A language and environment for statistical computing. R Foundation for Statistical Computing, Vienna, Austria

Sang T, Crawford D, Stuessy T (1997) Chloroplast DNA phylogeny, reticulate evolution, and biogeography of *Paeonia* (Paeoniaceae). Am J Bot 84:1120. doi: 10.2307/2446155

Sen S, Dayanandan S, Davis T, et al (2019) Origin and evolution of the genus Piper in Peninsular India. Mol Phylogenet Evol 138:102–113. doi: 10.1016/j.ympev.2019.05.033

Singh P, Karthigeyan K, Lakshminarasimhan P, Dash SS (2015) Endemic Vascular Plants of India. 1–339.

Stamatakis A (2014) RAxML version 8: a tool for phylogenetic analysis and post-analysis of large phylogenies. Bioinformatics 30:1312–1313. doi: 10.1093/bioinformatics/btu033

Suchard MA, Rambaut A (2009) Many-core algorithms for statistical phylogenetics. Bioinformatics 25:1370–1376. doi: 10.1093/bioinformatics/btp244

The Plant List (2013) The Plant List. In: The Plant List. 2013. Version 1.1. http://www.theplantlist.org/. Accessed 11 May 2020

Torres C (2000) Pollen size evolution: correlation between pollen volume and pistil length in Asteraceae. Sex Plant Reprod 12:365–370. doi: 10.1007/s004970000030

Tripp EA (2007) Evolutionary Relationships Within the Species-Rich Genus *Ruellia* (Acanthaceae). Syst Bot 32:628–649. doi: 10.1600/036364407782250625

Tripp EA (2010) Taxonomic Revision of *Ruellia* Section *Chiropterophila* (Acanthaceae): a Lineage of Rare and Endemic Species from Mexico. Syst Bot 35:629–661. doi: 10.1600/036364410792495845

Tripp EA, McDade LA (2014) A rich fossil record yields calibrated phylogeny for Acanthaceae (Lamiales) and evidence for marked biases in timing and directionality of intercontinental disjunctions. Syst Biol 63:660–684. doi: 10.1093/sysbio/syu029

Yu J, Xue J-H, Zhou S-L (2011) New universal *matK* primers for DNA barcoding angiosperms. J Syst Evol 49:176–181. doi: 10.1111/j.1759-6831.2011.00134.x

Zhisheng A, Kutzbach JE, Prell WL, Porter SC (2001) Evolution of Asian monsoons and phased uplift of the Himalaya-Tibetan plateau since Late Miocene times. Nature 411:62–66. doi: 10.1038/35075035

